# Potential host range of multiple SARS-like coronaviruses and an improved ACE2-Fc variant that is potent against both SARS-CoV-2 and SARS-CoV-1

**DOI:** 10.1101/2020.04.10.032342

**Authors:** Yujun Li, Haimin Wang, Xiaojuan Tang, Danting Ma, Chengzhi Du, Yifei Wang, Hong Pan, Qing Zou, Jie Zheng, Liangde Xu, Michael Farzan, Guocai Zhong

**Affiliations:** Shenzhen Bay Laboratory (SZBL), Shenzhen, China; School of Chemical Biology and Biotechnology, Peking University Shenzhen Graduate School, Shenzhen, China; Shanghai Institute of Materia Medica, Chinese Academy of Sciences, Shanghai, China; School of Biomedical Engineering and Eye Hospital, Wenzhou Medical University, Wenzhou, China; Department of Immunology and Microbiology, The Scripps Research Institute, Jupiter, Florida, United States

## Abstract

The severe acute respiratory syndrome coronavirus 2 (SARS-CoV-2) is a currently uncontrolled pandemic and the etiological agent of coronavirus disease 2019 (COVID-19). It is important to study the host range of SARS-CoV-2 because some domestic species might harbor the virus and transmit it back to humans. In addition, insight into the ability of SARS-CoV-2 and SARS-like viruses to utilize animal orthologs of the SARS-CoV-2 receptor ACE2 might provide structural insight into improving ACE2-based viral entry inhibitors. Here we show that ACE2 orthologs of a wide range of domestic and wild animals support entry of SARS-CoV-2, as well as that of SARS-CoV-1, bat coronavirus RaTG13, and a coronavirus isolated from pangolins. Some of these species, including camels, cattle, horses, goats, sheep, pigs, cats, and rabbits may serve as potential intermediate hosts for new human transmission, and rabbits in particular may serve as a useful experimental model of COVID-19. We show that SARS-CoV-2 and SARS-CoV-1 entry could be potently blocked by recombinant IgG Fc-fusion proteins of viral spike protein receptor-binding domains (RBD-Fc) and soluble ACE2 (ACE2-Fc). Moreover, an ACE2-Fc variant, which carries a D30E mutation and has ACE2 truncated at its residue 740 but not 615, outperforms all the other ACE2-Fc variants on blocking entry of both viruses. Our data suggest that RBD-Fc and ACE2-Fc could be used to treat and prevent infection of SARS-CoV-2 and any new viral variants that emerge over the course of the pandemic.

## INTRODUCTION

Two subgroups of coronaviruses, including alphacoronaviruses (e.g. swine acute diarrhea syndrome coronavirus, SADS-CoV) and betacoronaviruses (e.g. SARS coronavirus 1, SARS-CoV-1), infect mammals and have broad host ranges spanning bats, rodents, domestic animals, and humans^1–7^. Bats are considered to be the natural reservoir hosts for a number of pathogenic human coronaviruses, including SARS-CoV-1, Middle East respiratory syndrome coronavirus (MERS-CoV), HCoV-NL63, and HCoV-229E ^6,8^. Domestic animals have been shown to play key roles as intermediate hosts, such as camels for MERS-CoV and camelids for HCoV-229E, to transmit coronaviruses from bats to humans ^6^. The severe acute respiratory syndrome coronavirus 2 (SARS-CoV-2), the etiological agent of coronavirus disease 2019 (COVID-19)^8–10^, shares 79.5% genome sequence identity to SARS-CoV-1 and 96.2% genome sequence identity to a bat coronavirus, Bat-CoV RaTG13, indicating that bats are also likely the reservoir hosts for SARS-CoV-2 ^ref.8–10^. However, it is unknown whether domestic animals also served as intermediate hosts that facilitated SARS-CoV-2 transmission from its natural reservoir hosts to humans.

As of April 7^th^ 2020, the uncontrolled COVID-19 pandemic has already caused over 1,210,000 confirmed infections and over 67,000 documented deaths across 211 countries, according to World Health Organization’s online updates. There are currently no vaccines or targeted therapeutics available for this disease^11^. It is therefore urgent to study the host range of SARS-CoV-2 because some domestic animals may harbor and spread the virus. More importantly, studies on the host range of SARS-CoV-2 may also help identify new model animals to facilitate basic and translational studies on COVID-19.

Receptor binding is a critical determinant for the host range of coronaviruses^6,12^. SARS-CoV-2 utilizes ACE2, the SARS-CoV-1 receptor^13^, as an essential cellular receptor to infect cells^8,14,15^. Therefore, in this study, we characterized seventeen ACE2 orthologs for their function to support entry of SARS-CoV-2, as well as that of SARS-CoV-1, a bat coronavirus (Bat-CoV RaTG13)^ref.8^, and a coronavirus recently isolated from Malayan pangolins (called Pangolin-CoV hereafter)^16^. Furthermore, to explore broadly anti-coronavirus approaches to treatment and prevention of SARS-CoV-2, we evaluated recombinant receptor-binding domain (RBD) variants of the viral spike proteins, and soluble ACE2 variants for their potency as entry inhibitors against SARS-CoV-2 and SARS-CoV-1.

## RESULTS

### A wide range of ACE2 orthologs support binding to RBD proteins of SARS-CoV-2, Pangolin-CoV, Bat-CoV RaTG13, and SARS-CoV-1

The RBD of SARS-CoV-2, like that of SARS-CoV-1, mainly relies on residues in its receptor binding motif (RBM) to bind ACE2 (**Figs. 1a** and **b**)^17–19^. Through analyzing ACE2 residues that are within 5 Å from SARS-CoV-2 RBD atoms, we found that these residues are highly conserved across the seventeen analyzed ACE2 orthologs, including human ACE2 and ACE2 orthologs of sixteen domestic and wild animals (**Figs. 1a** and **c**, **Extended Data Fig. 1**, and **Extended Data Table. 1**). We then made expression plasmids for the seventeen ACE2 orthologs and expressed mouse IgG2 Fc fusion proteins of the RBDs of four coronaviruses, including SARS-CoV-2 WHU01, Bat-CoV RaTG13, Pangolin-CoV, and SARS-CoV-1 BJ01 (**Extended Data Fig. 2a**). These RBD proteins were then used to perform surface staining of 293T cells transfected with each of the seventeen ACE2 orthologs or a vector plasmid control (**Figs. 2a** and **b**). The flow cytometry data clearly show that, although each of the four tested RBDs has a distinctive binding profile, they all can bind to a wide range of ACE2 orthologs. Note that human ACE2 and ACE2 orthologs of all the domestic animals, except for that of donkeys, guinea pigs, and chickens, support efficient binding to all the four tested RBDs. The lack of any RBD binding to donkey ACE2 is likely due to a protein sequence defect, because expression of the donkey ACE2 is detectable in 293T cells but a 22-residue fragment likely important for ACE2 folding is missing in the Genbank sequence (**Fig. 2c**, and **Extended Data Figs. 1** and **2b**). These data suggest that ACE2 orthologs of domestic animals might be generally functional for supporting cell entry of the four tested viruses.

**Fig. 1.**
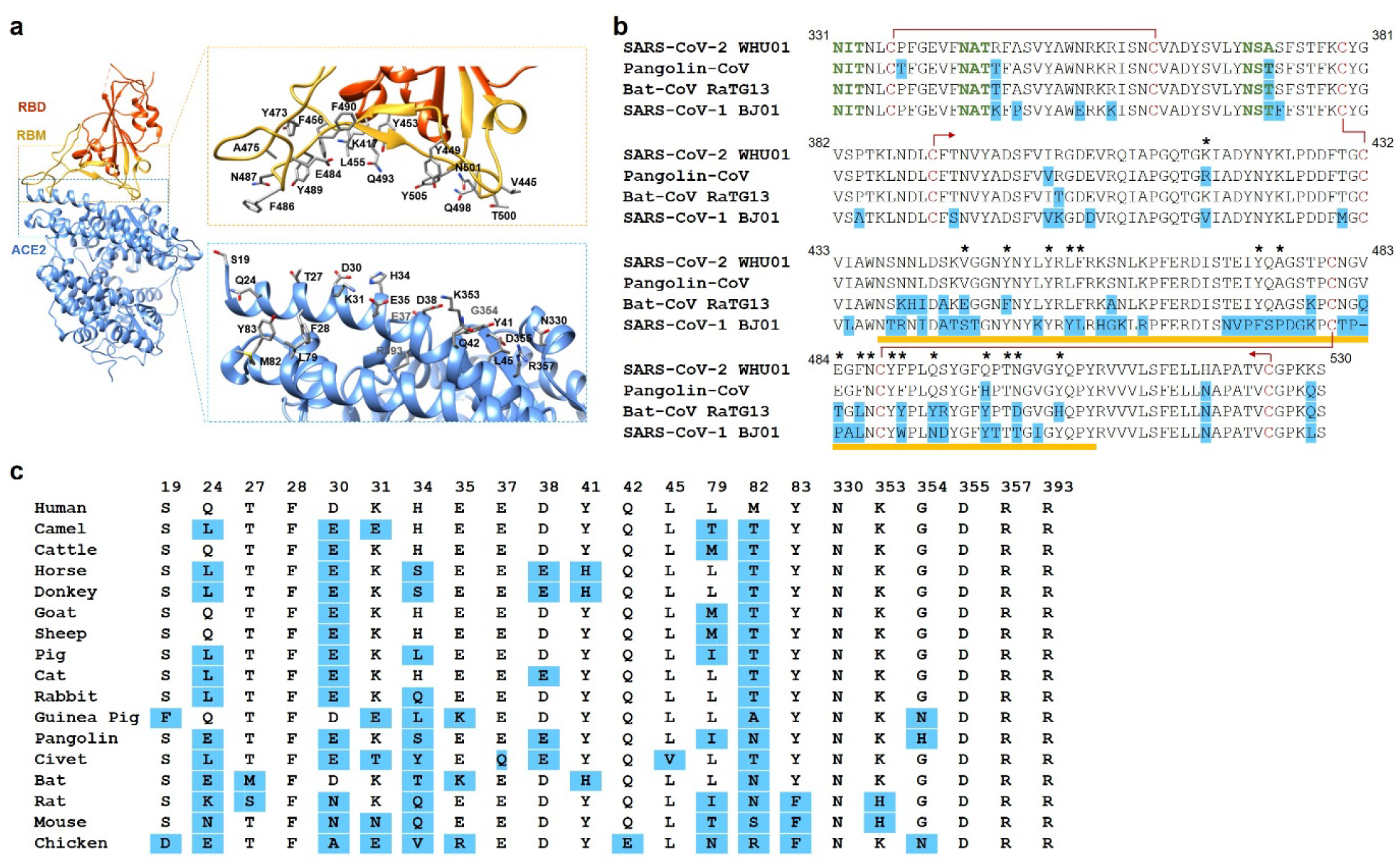
SARS-CoV-2 and ACE2 contact residues are conserved among SARS-like viruses and ACE2 orthologs, respectively. **a**, Interactions between the SARS-CoV-2 receptor binding domain (RBD, red) and ACE2 (blue) involve a large number of contact residues (PDB code 6M0J). RBD residues in less than 5 Å from ACE2 atoms and ACE2 residues in less than 5 Å from RBD atoms are shown. **b**, The sequences of the SARS-CoV-2 WHU01, a pangolin coronavirus (Pangolin-CoV), a bat coronavirus (Bat-CoV RaTG13), and the SARS-CoV-1 BJ01 are aligned, with residues different from the corresponding ones in SARS-CoV-2 highlighted in blue. The stars indicate RBD residues in less than 5 Å from ACE2 atoms. The yellow lines indicate the RBM region, and the red lines or arrows indicate disulfide linkages. N-linked glycosylation motifs are indicated in green. **c**, Sequences of ACE2 orthologs from the 17 indicated species are aligned, with only residues in less than 5 Å from RBD atoms shown here and aligned full-length protein sequences shown in the **Extended Data Figure 1**. The numbering is based on human ACE2 protein, and the residues different from the corresponding ones in human ACE2 are highlighted in blue.

**Fig. 2.**
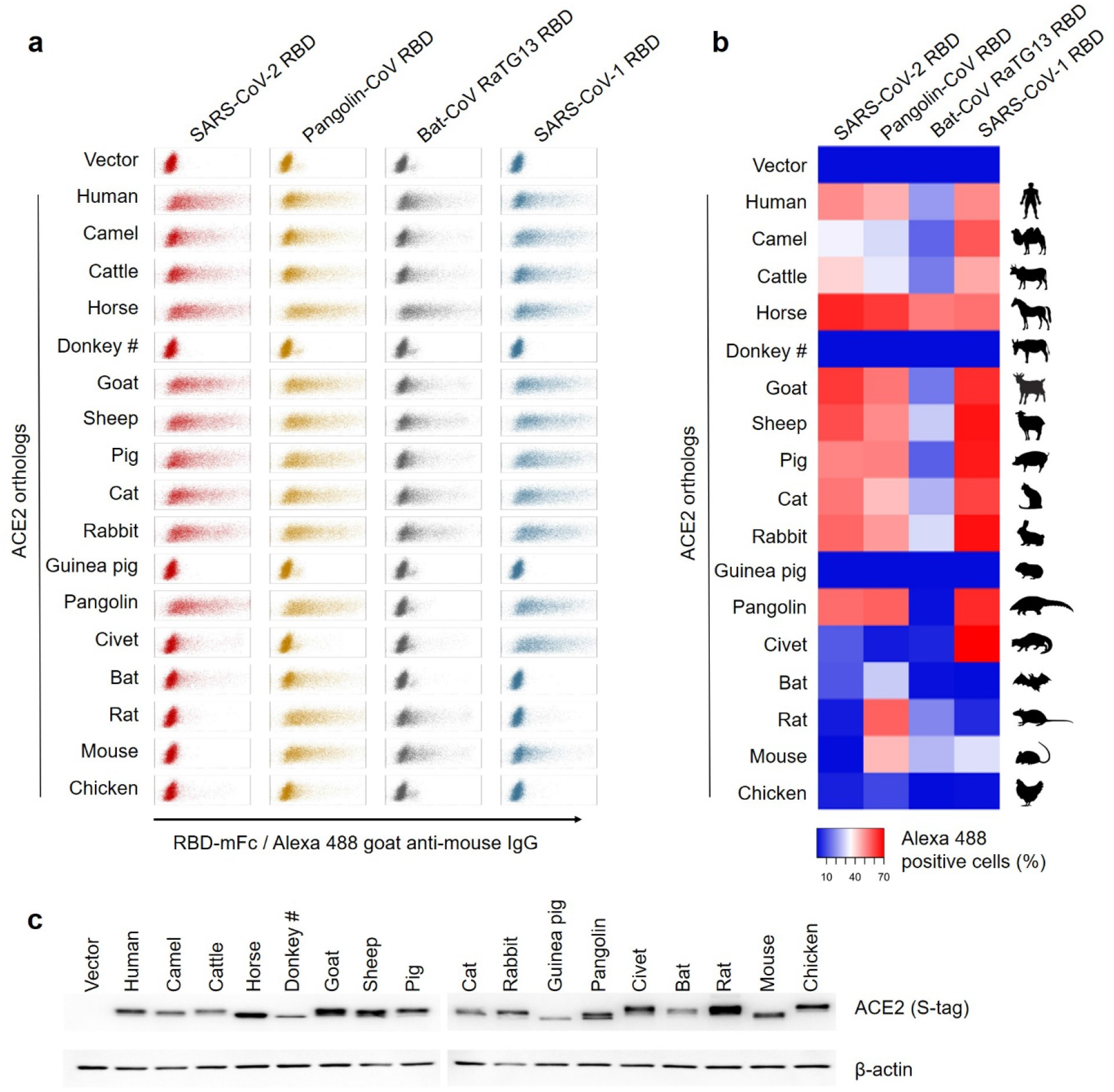
A wide range of ACE2 orthologs support binding to RBD proteins of four SARS-like coronaviruses. **a**, 293T cells transfected with ACE2 genes of the indicated species were stained with RBD-mouse IgG2 Fc fusion proteins of SARS-CoV-2 WHU01, Pangolin-CoV, Bat-CoV RaTG13, and SARS-CoV-1 BJ01. The cells were then stained with Alexa 488 goat anti-mouse IgG secondary antibody and RBD-ACE2 binding was detected using flow cytometry. #, based on ACE2 protein sequence alignment shown in the Extended Data Figure 1, around 20 residues (246-268) of donkey ACE2 are missing in the sequence used in this study (NCBI Reference Sequence: XM_014857647.1). **b**, Percentages of cells positive for RBD binding in panel *a* are presented as a heatmap according to the indicated color code. **c**, Expression levels of the indicated ACE2 genes were detected using Western Blot. Data shown are representative of two independent experiments with similar results.

### A wide range of ACE2 orthologs, in particular that of rabbits, support entry of the four coronaviruses

To evaluate spike protein-mediated entry of these coronaviruses, we first tested multiple conditions for establishing a robust SARS-CoV-2 pseudovirus reporter assay (**Figs. 3a** and **b**, and **Extended Data Fig. 3a**). We found that, although the transmembrane serine protease TMPRSS2 slightly enhanced the pseudovirus-produced reporter signals under certain conditions, the most robust signals were produced by the reporter retroviruses pseudotyped with a SARS-CoV-2 spike variant that had the ‘PRRAR’ furin site mutated to a single arginine residue. Then reporter retroviruses pseudotyped with spike proteins of SARS-CoV-2, its furin site deletion mutant (ΔFurin), Bat-CoV RaTG13, and SARS-CoV-1, were respectively used to infect 293T cells expressing each of the seventeen ACE2 orthologs. A VSV-G-pseudotyped reporter retrovirus whose entry is independent of ACE2 was used as a control virus. Consistent with the binding data, the infection data clearly show that SARS-CoV-2, SARS-CoV-1, and Bat-CoV RaTG13 have distinctive infection profiles and ACE2 orthologs of most domestic animals can support entry of all three viruses (**Fig. 3c**, and **Extended Data Fig. 3b**). These data suggest that domestic animals, including camels, cattle, horses, goats, sheep, pigs, cats, and rabbits may serve as potential intermediate hosts for new human transmission. Moreover, the finding that rabbit ACE2 efficiently supports entry of all the three tested viruses suggests that rabbits may serve as a useful model animal of COVID-19, as well as that of SARS and diseases caused by other SARS-related coronaviruses.

**Fig. 3.**
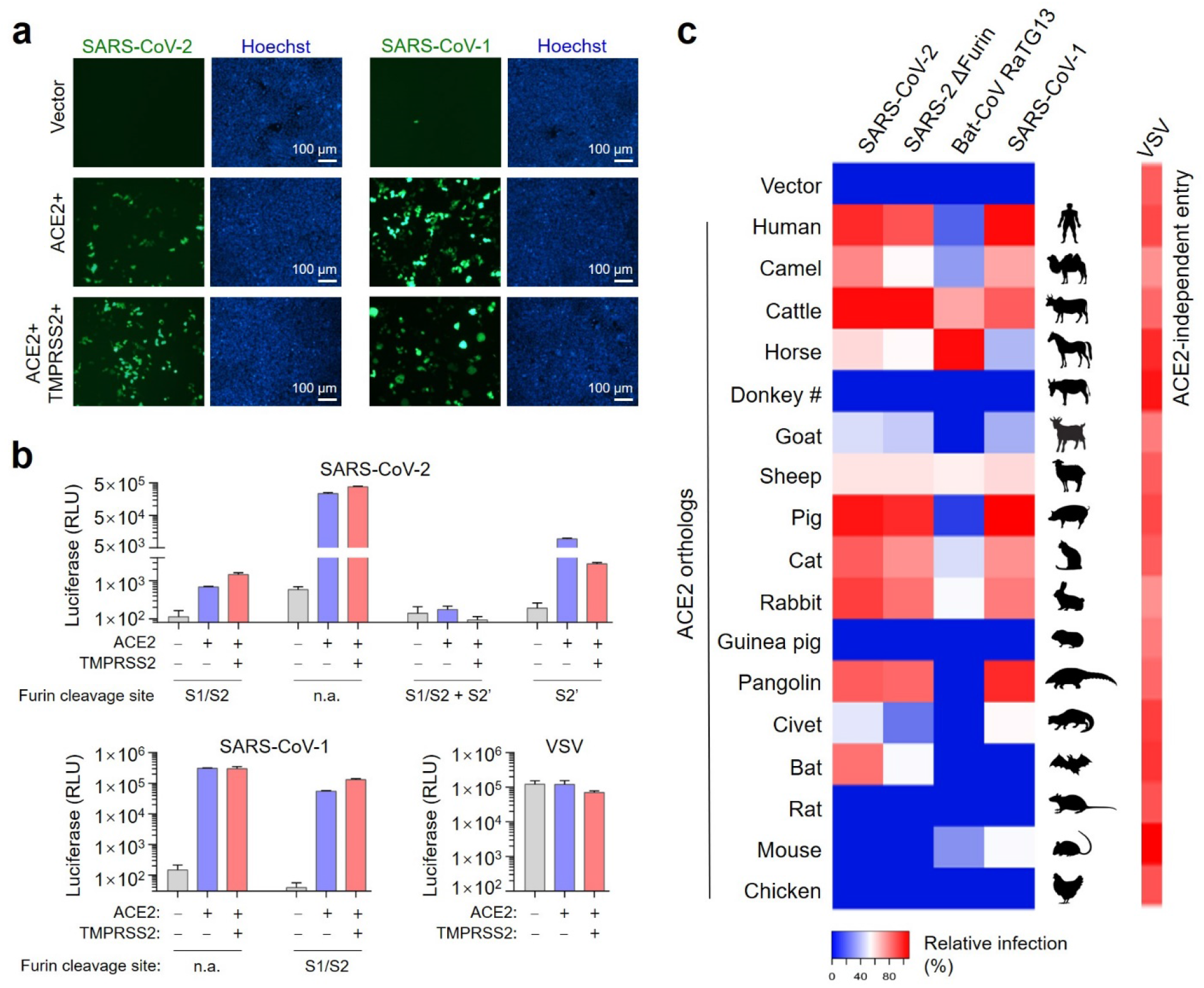
A wide range of ACE2 orthologs support entry of SARS-CoV-2, Bat-CoV RaTG13, and SARS-CoV-1. **a**, 293T cells were transfected with a vector control plasmid, an ACE2 expression plasmid, or ACE2 and TMPRSS2 expression plasmids. The cells were then infected with GFP reporter retroviruses pseudotyped with SARS-CoV-2 and SARS-CoV-1 spike proteins, respectively. Spike protein-mediated coronavirus entry was detected by fluorescent microscopy at 48 hours post infection. **b**, Experiments similar to *a* except that retroviruses carrying a luciferase reporter gene and pseudotyped with SARS-CoV-2, SARS-CoV-1 spike mutants and VSV-G were used. Coronavirus spike mutant-mediated viral entry was measured by the luciferase reporter at 48 hours post infection. **c**, 293T cells expressing the indicated ACE2 orthologs were infected with luciferase-reporter retroviruses pseudotyped with the indicated spike proteins. ACE2 ortholog-mediated viral entry was measured by the luciferase reporter at 48 hours post infection. Original reporter expression data are provided in **Extended Data Figure 3b**. Relative infection (%) values for each ACE2 ortholog-mediated viral entry were independently calculated against the highest expression values of the same pseudotype panel and presented as a heatmap according to the indicated color code. Data shown are representative of three experiments independently performed by three different people with similar results, and data points represent mean ± s.d. of three biological replicates.

### Recombinant RBD-Fc and ACE2-Fc proteins potently block entry of SARS-CoV-2 and SARS-CoV-1

Recombinant RBD and soluble ACE2 proteins have been shown to potently block SARS-CoV-1 entry^20,21^. To investigate whether similar approaches could also be applied to SARS-CoV-2, we first produced IgG Fc fusion proteins of RBD (RBD-Fc) and soluble ACE2 (ACE2-Fc) variants (**Fig. 4a**). Specifically, the RBD variants include wildtype RBDs of SARS-CoV-1, Pangolin-CoV, and SARS-CoV-2, and four RBD mutants of SARS-CoV-2 (**Extended Data Figs. 2a** and **4a**). The ACE2-Fc variants include human ACE2 truncated at its residue 615 (615-wt) and 740 (740-wt), respectively, and mutants of the 615- and 740-version ACE2-Fc proteins (**Extended Data Fig. 4b**). We then evaluated these proteins for their potency of blocking SARS-CoV-2 pseudovirus infection. Among all the RBD-Fc variants, the Y505W mutant of SARS-CoV-2 RBD showed modestly improved potency over the wildtype, and wildtype RBD of Pangolin-CoV showed the best neutralization activity among all the tested RBDs (**Fig. 4b**). Among the tested ACE2-Fc variants, interestingly, all the 740-version variants showed significantly better potency than the 615-version variants (two-tailed two-sample t-test, p<0.001; **Fig. 4c**). In addition, D30E mutants of both 615- and 740-version proteins showed improved potency over the corresponding wildtypes (**Fig. 4c**). We further tested three of the RBD-Fc variants and four of the ACE2-Fc variants for their neutralization activities against SARS-CoV-1 and observed similar results, except that D30E mutation did not show any improvement on the neutralization potency against this virus (**Figs. 4d** and **e**). Consistently, cell-surface ACE2 carrying this D30E mutation binds SARS-CoV-2 RBD better than the wildtype ACE2 does, and this enhanced binding was not observed with SARS-CoV-1 RBD (**Extended Data Fig. 4c**). Thus, the mechanism of the D30E-mediated improvement is likely that the mutation enhances the salt-bridge interaction between the residue 30 of the ACE2 and residue 417 of the SARS-CoV-2 RBD (**Fig. 4f**), because this interaction is only present in the ACE2 complex with the RBD of SARS-CoV-2 but not that of SARS-CoV-1 ^ref.17–19^.

**Fig. 4.**
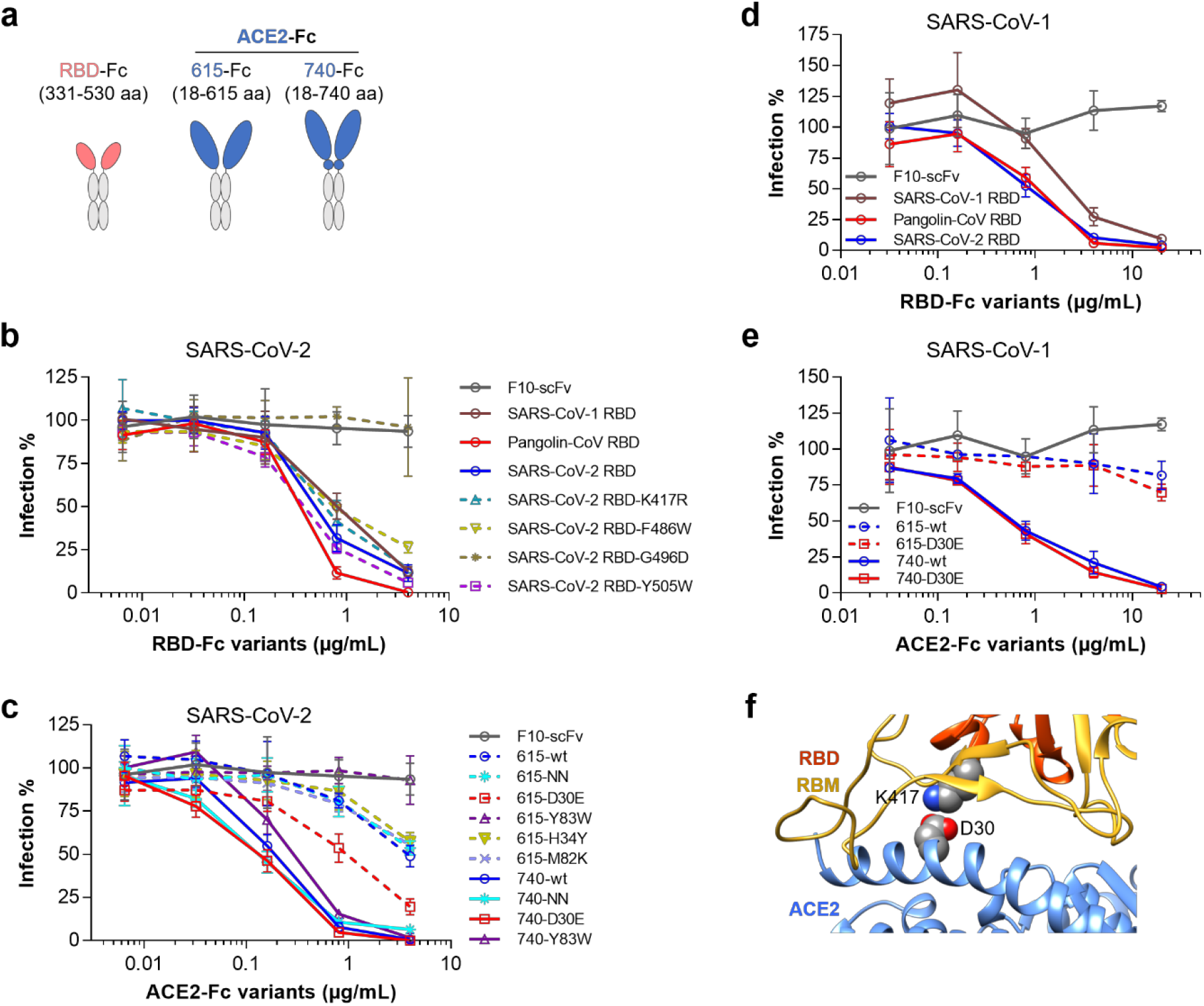
Recombinant RBD-Fc and ACE2-Fc variants efficiently block entry of SARS-CoV-2 and SARS-Cov-1. **a**, Diagrams of RBD-Fc and ACE2-Fc fusion proteins used in the following studies. ACE2-expressing 293T cells were infected with SARS-CoV-2 spike-pseudotyped retrovirus in the presence of purified recombinant RBD-Fc (**b**) and ACE2-Fc (**c**) fusion proteins at the indicated concentrations. Viral entry was measured by the luciferase reporter at 48 hours post infection. Note that all the 740-version variants showed significantly better potency than the 615-version variants (two-tailed two-sample t-test, p<0.001). **d-e**, Experiments similar to that of panels b and c, except that the indicated RBD-Fc (d) and ACE2-Fc (e) variants were tested against SARS-CoV-1 pseudovirus. **f**, The ACE2 residue D30 forms a salt bridge with the SARS-CoV-2 RBD residue K417 (PDB code 6M0J). Data shown are representative of three experiments independently performed by three different people with similar results, and data points represent mean ± s.d. of four biological replicates.

## DISCUSSION

In this study, we found that ACE2 orthologs of a wide range of domestic animals and pangolins efficiently support binding and entry of SARS-CoV-2, as well as that of SARS-CoV-1, a pangolin coronavirus (Pangolin-CoV), and bat coronavirus RaTG13. In addition, the pangolin coronavirus tested here showed the broadest host range. Our data therefore suggest that livestock and companion pets, including camels, cattle, horses, goats, sheep, pigs, cats, and rabbits are potential intermediate hosts for these viruses. Therefore, to prevent new outbreaks of SARS-CoV-2 transmission, it is necessary to consider epidemic surveillance of livestock animals in marketplaces and companion pets from families that have COVID-19 patients.

Animal models are essential for pre-clinical evaluation of efficacy and potential toxicity of candidate prophylactic vaccines or therapeutics for COVID-19. They are also very useful for studying transmission, pathogenesis, and immunology of this disease. Human *ACE2* transgenic mice have been shown to support SARS-CoV-2 replication in the lung, lose weight early after infection and then develop anti-spike IgG immune responses^22^. However, the expression patterns of a constitutively expressed *ACE2* transgene may not fully follow the patterns of endogenous *ACE2* gene. There are also major differences between mice and humans in immune responses against pathogens, with immune resistance dominant in humans but tolerance mechanisms dominant in mice^23^. Therefore, it’s necessary to find better model animals, which can use endogenous ACE2 to support SARS-CoV-2 infection and whose immune systems better mimic human’s. Our finding that rabbit ACE2 efficiently supports binding and entry of the tested diverse coronaviruses suggests that rabbits may serve as a new model animal of COVID-19, as well as that of SARS and diseases caused by other SARS-related coronaviruses.

Although in urgent needs, vaccines and targeted therapeutics for SARS-CoV-2 infections are not available yet^11^. More importantly, anti-spike IgG, either passively administered or vaccine-induced, was also found to cause severe acute lung injury during acute SARS-CoV infection, likely through an antibody-dependent enhancement mechanism^24,25^. Thus, in addition to pursuing traditional vaccination strategies and isolation of fully human anti-spike IgG antibodies from convalescent-phase patient peripheral blood mononuclear cells (PBMCs), alternative approaches to prevention and treatment of SARS-CoV-2 infection should also be explored. Here we found that four diverse SARS-like coronaviruses can efficiently utilize ACE2 for entry, and that both RBD-Fc and ACE2-Fc proteins, in particular the 740-version of ACE2-Fc and its D30E mutant, potently blocked entry of both SARS-CoV-2 and SARS-CoV-1. These data provide very encouraging evidence that these proteins might be developed as effective prophylactic and therapeutic agents against infection of SARS-CoV-2 and any new viral variants that emerge over the course of the pandemic.

## METHODS

Methods and associated references are provided below.

## ACKNOWLEGEMENTS

This work is supported by Shenzhen Bay Laboratory Startup Funds No.21230041. We thank Dr. Yu J. Cao (School of Chemical Biology and Biotechnology, Peking University Shenzhen Graduate School, Shenzhen China) for generously providing the 293F cells used for recombinant protein production in this study.

## CONTRIBUTIONS

G.Z. conceived and designed this study. Y.L., H.W., X.T., D.M., C.D., Y.W., H.P., Q.Z., and G.Z. performed all experiments, acquired and analyzed all data. J.Z., and L.X. contributed key reagents. M.F. provided key insights that helped the design of this study, analyzed the data, and revised the manuscript. G.Z. wrote the manuscript.

## COMPETING INTERESTS

The authors declare that no competing interest exists.

## METHODS

### Plasmids

DNA fragments encoding C-terminally S-tagged ACE2 orthologs were synthesized in pUC57 backbone plasmid by Sangon Biotech (Shanghai, China). These fragments were then cloned into pQCXIP plasmid (Clontech) between *Sbf*I and *Not*I restriction sites. DNA fragments encoding spike proteins of SARS-CoV-2 WHU01 (GenBank: MN988668.1), SARS-CoV-1 BJ01 (GenBank: AY278488.2), Pangolin-CoV (National Genomics Data Center: GWHABKW00000000)^1^, and Bat-CoV RaTG13 (GenBank: MN996532)^2^, were synthesized by the Beijing Genomic Institute (BGI, China) and Sangon Biotech (Shanghai, China) and then cloned into pcDNA3.1(+) plasmid or pCAGGS plasmid between *Eco*RI and *Xho*I restriction sites. Plasmids encoding recombinant RBD and soluble ACE2 variants were generated by cloning each of the gene fragments into a pCAGGS-based mouse IgG2a Fc fusion protein expression plasmid between *Not*I and *Bsp*EI sites. The retroviral reporter plasmids encoding a *Gaussia* luciferase or a green fluorescent protein (GFP) reporter gene were constructed by cloning the reporter genes into pQCXIP plasmid (Clontech), respectively.

### Cell culture

293T cell line was purchased from Stem Cell Bank, Chinese Academy of Sciences, and confirmed mycoplasma-free by the provider. 293T cells were maintained in Dulbecco’s Modified Eagle Medium (DMEM, Life Technologies) at 37°C in a 5% CO_2_-humidified incubator. Growth medium were supplemented with 2 mM Glutamax-I^™^ (Gibco), 100 μM non-essential amino acids (Gibco), 100 U/mL penicillin and 100 μg/mL streptomycin (Gibco), and 10% FBS (Gibco). 293T-based stable cells expressing human ACE2 were maintained under the same culture condition as 293T, except that 3 μg/mL of puromycin was added to the growth medium. 293F cells for recombinant protein production was generously provided by Dr. Yu J. Cao (School of Chemical Biology and Biotechnology, Peking University Shenzhen Graduate School) and maintained in SMM 293-TII serum-free medium (Sino Biological, China) at 37 °C, 8% CO2, in a shaker incubator at 125 rpm.

### IgG Fc fusion protein production and purification

293F cells at the density of 6 × 10^5^ cells/mL were seeded into 100 mL SMM 293-TII serum-free medium (Sino Biological, China) one day before transfection. Cells were then transfected with 100 μg plasmid in complex with 250 μg PEI MAX 4000 (Ploysciences, Inc, USA). Cell culture supernatants were collected at 48 to 72 hours post transfection. Recombinant Fc fusion proteins are purified using Protein A Sepharose CL-4B (GE Healthcare), eluted with 0.1 M citric acid at pH 4.5 and neutralized with 1 M Tris-HCl at pH 9.0. Buffers were then exchanged to PBS and proteins were concentrated by 30 kDa cut-off Amicon Ultra-15 Centrifugal Filter Units (Millipore).

### Flow cytometry for detecting interactions of RBD-Fc proteins with cell surface ACE2 orthologs

293T cells were seeded at 20% density in 48-well plates at 12-15 hours before transfection. Cells in each well were then transfected with 0.5 μL of lipofectamine 2000 (Life Technologies) in complex with 200 ng of plasmid encoding one of the seventeen ACE2 orthologs or a D30E mutant of the human ACE2. Culture medium was changed at 6 hours after transfection. Cells were then detached with 5 mM EDTA at 36 hours post transfection. Cells were then stained with 5 μg/mL RBD-Fc proteins at 37 °C for 10 min, washed three times, and then stained with 2 μg/mL Alexa488 conjugated goat anti-mouse IgG secondary antibody (Invitrogen, Cat. No. A-11001) at room temperature for 20 min. After another three washes, cells were analyzed by Attune NxT flow cytometer (Thermo Fisher) and signals of 10,000 FSC/SSC gated cells were collected for each sample.

### Western Blot to detect S-tagged ACE2 (ACE2-S-tag) or C9-tagged spike (Spike-C9-tag) expression in 293T cells

293T cells were seeded at 20% density in 6-well plates at 12-15 hours before transfection. Cells in each well were then transfected with 2 μg of plasmid in complex with 5 μL of lipofectamine 2000 (Life Technologies). Thirty-six hours after transfection, cells were lysed and 10 μg of total protein were used for western blot. ACE2-S-tag expression was detected by using 6.2, a mouse anti-S-tag monoclonoal antibody (Invitrogen, Cat. No. MA1-981), and an HRP-conjugated goat anti-mouse IgG Fc secondary antibody (Invitrogen, Cat. No. 31437). **Spike-C9-tag** expression was then detected by using 1D4, a mouse anti-C9-tag monoclonal antibody (Invitrogen, Cat. No. MA1-722), and the HRP-conjugated goat anti-mouse IgG Fc secondary antibody (Invitrogen, Cat. No. 31437).

### Production of reporter retroviruses pseudotyped with coronavirus spike proteins or VSV-G

293T cells were seeded at 30% density in 150 mm dish at 12-15 hours before transfection. Cells were then transfected with 67.5 μg of polyethylenimine (PEI) Max 40,000 (Polysciences) in complex with 11.25 μg of plasmid encoding a coronavirus spike protein or VSV-G, 11.25 μg of plasmid encoding murine leukemia virus (MLV) Gag and Pol proteins, and 11.25 μg of a pQCXIP-based GFP or luciferase reporter plasmid. Eight hours after transfection, cell culture medium was refreshed and changed to growth medium containing 2% FBS (Gibco) and 25 mM HEPES (Gibco). Cell culture supernatants were collected at 36-48 hours post transfection, spun down at 3000×g for 10 min, and filtered through 0.45 μm filter units to remove cell debris. Coronavirus spike-pseudotyped viruses were then concentrated 10 times at 2000×g using 100 kDa cut-off Amicon Ultra-15 Centrifugal Filter Units (Millipore).

### Pseudovirus infection of 293T cells expressing ACE2 orthologs

293T cells were seeded at 20% density in poly-lysine pre-coated 48-well plates 12-15 hours before transfection. Cells in each well were then transfected with 0.5 μL of lipofectamine 2000 (Life Technologies) in complex with 200 ng of a vector control plasmid or a plasmid encoding one of the seventeen ACE2 orthologs. Cell culture medium was refreshed at 6 hours post transfection. Additional 18 hours later, cells in each well were infected with 50 μL of SARS-CoV-2 wildtype pseudovirus (10×concentrated), 20 μL of SARS-CoV-2 ΔFurin pseudovirus (10×concentrated), 50 μL of Bat-CoV RaTG13 pseudovirus (10×concentrated), 10 μL of SARS-CoV-1 pseudovirus (10×concentrated), or 10 μL of VSV-G pseudovirus diluted in 200 μL of culture medium containing 2% FBS (Gibco). Culture medium was refreshed at 2 hours post pseudovirus infection and the medium was refreshed every 12 hours. For the luciferase reporter virus-infected cells, cell culture supernatants were collected and subjected to a *Gaussia* luciferase assay at 48 hours post infection. The GFP reporter virus-infected cells were stained with Hoechst 33342 (Invitrogen) and subjected to fluorescent microscopy (IX73 microscope, Olympus) at 48 hours post infection.

### *Gaussia* luciferase luminescence flash assay

To measure *Gaussia* luciferase expression, 20 μL of cell culture supernatant of each sample and 100 μL of assay buffer containing 4 μM coelenterazine native (Biosynth Chemistry & Biology) were added to one well of a 96-well black opaque assay plate (Corning), and measured with the Tristar5 multifunctional microplate reader (Berthold Technologies) for 0.1 second/well.

### Coronavirus pseudovirus neutralization assay

293T cells stably expressing human ACE2 protein were seeded at 20% density in poly-lysine pre-coated 48-well plates 18-24 hours before pseudovirus infection. SARS-CoV-2 or SARS-CoV-1 spike protein-pseudotyped luciferase reporter virus was diluted in DMEM (2% FBS, heat-inactivated) containing titrated amounts of RBD-Fc or ACE2-Fc variant proteins. Virus-inhibitor mixtures were then added to cells for 2 h at 37 °C. Cells were then washed with serum-free medium and incubated in 250 μL of DMEM (2% FBS) at 37 °C. Cell culture supernatants were collected for the Gaussia luciferase assay at 48 h post infection.

### Statistical analysis

Data expressed as mean values ± s.d. or s.e.m. Statistical analyses were performed using two-sided two-sample Student’s *t*-test using GraphPad Prism 6.0 software when applicable. Differences were considered significant at *P* < 0.01.

## Data availability

Our research resources, including methods, plasmids, and protocols, are available upon reasonable request to qualified academic investigators for noncommercial research purposes. All reagents developed, such as vector plasmids, as well as detailed methods, will be made available upon written request.

## EXTENDED DATA FIGURES 1-4 AND TABLE 1

**Extended Data Fig. 1.**
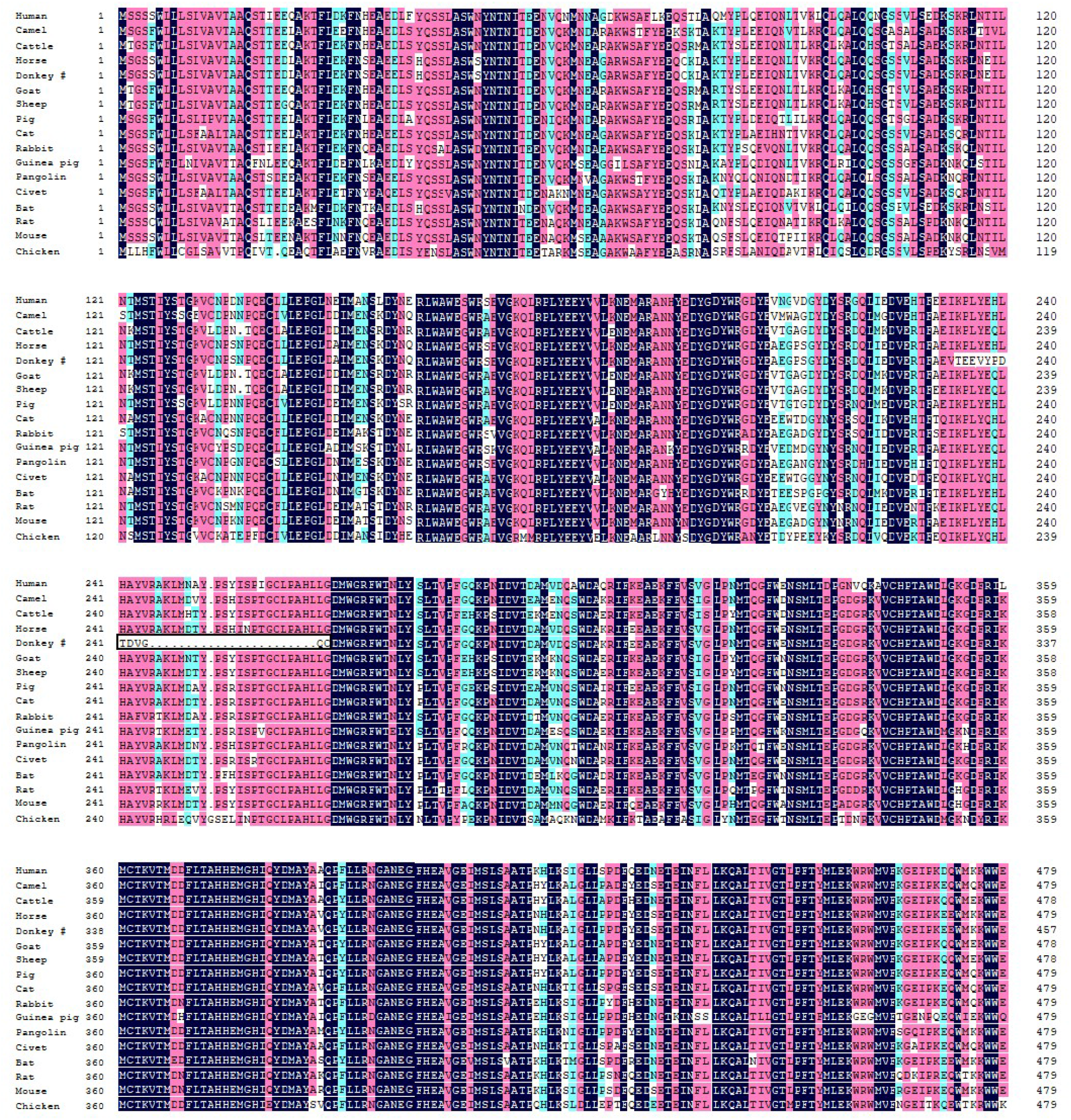
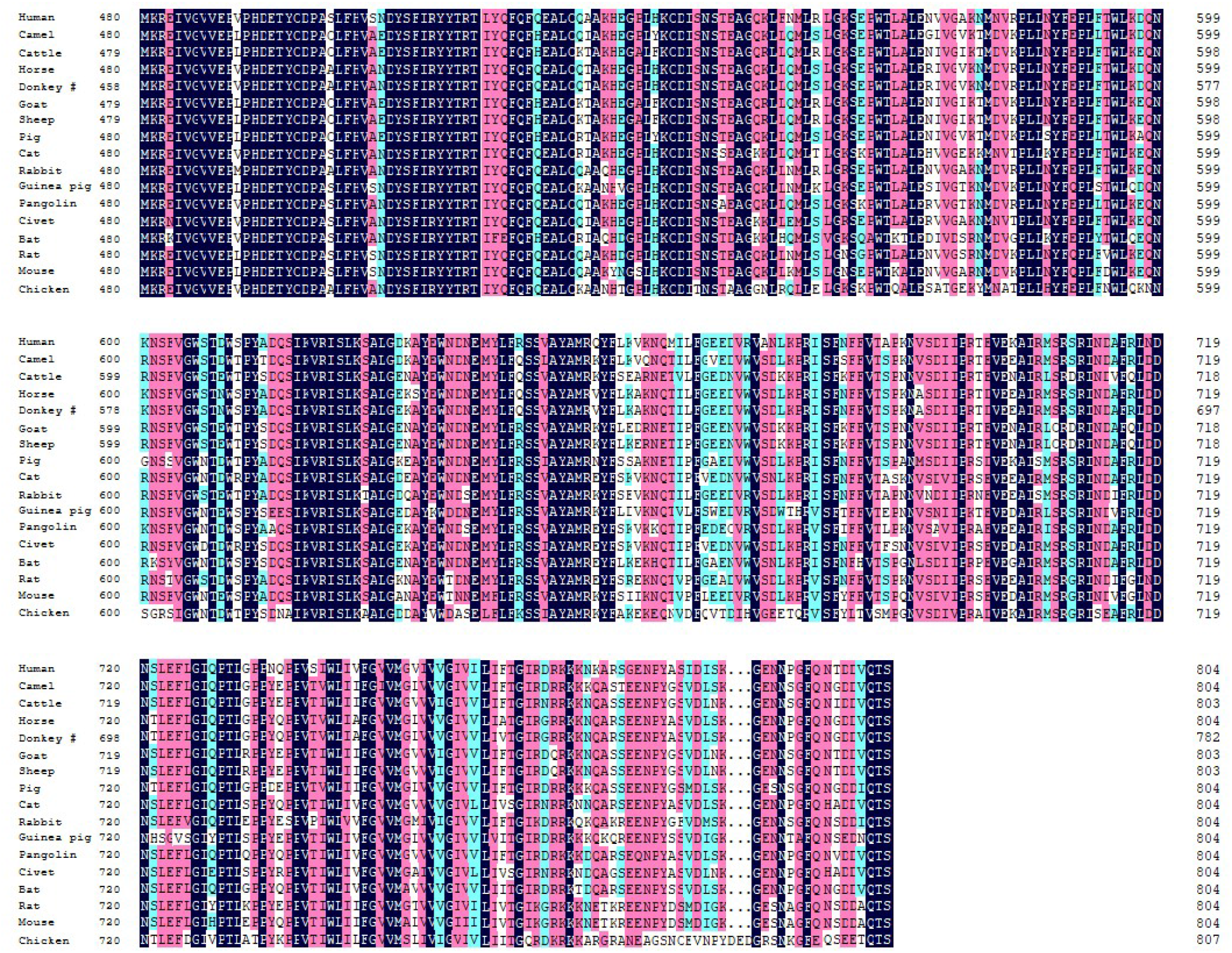
Protein sequence alignment of the seventeen ACE2 orthologs evaluated in this study. Detailed information about the species and accession numbers of the ACE2 orthologs are presented in the Extended Data Table 1.

**Extended Data Fig. 2.**
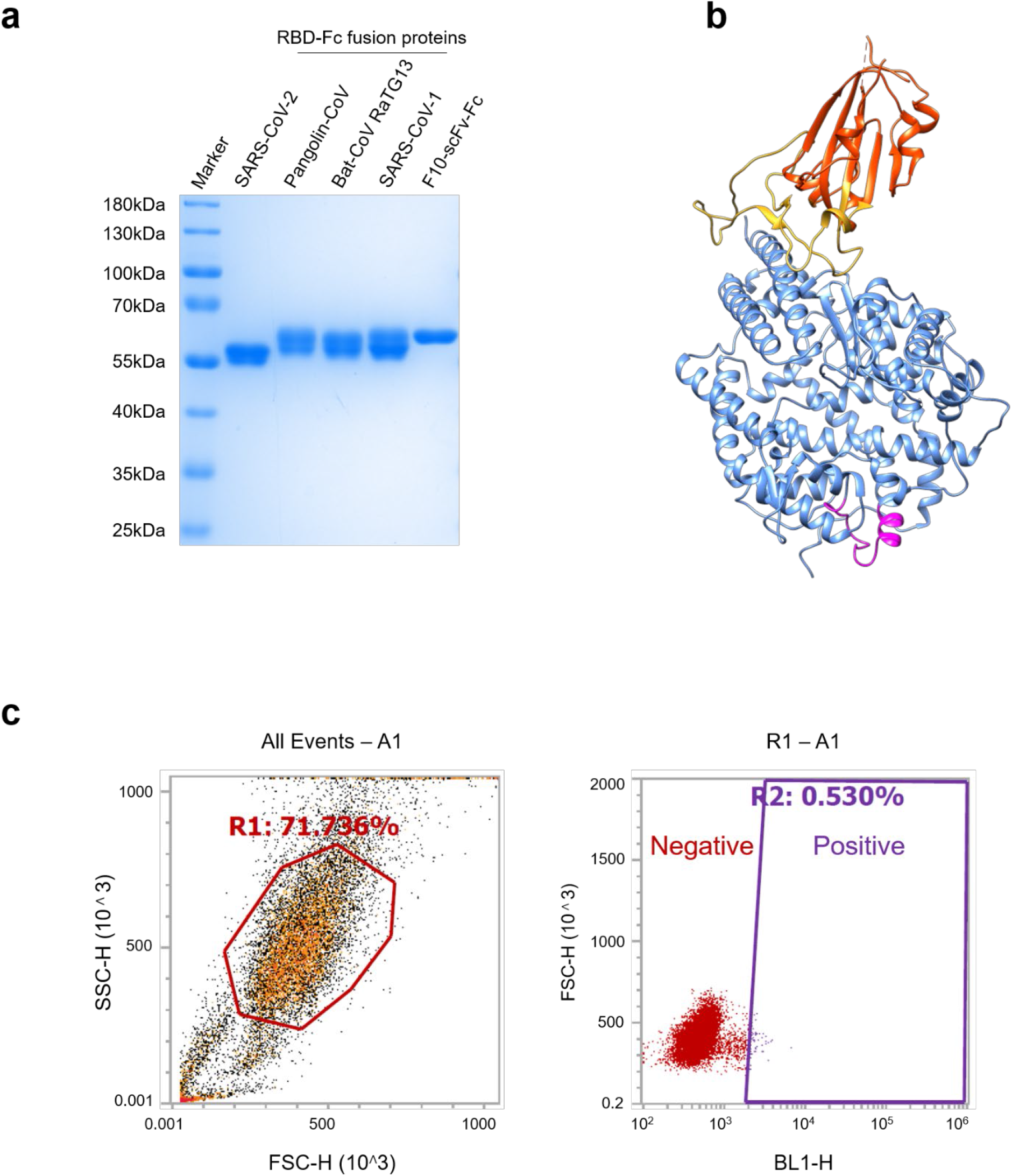
Key reagents used in the experiments shown in Figure 2. **a**, SDS-PAGE image of the purified recombinant RBD-Fc fusion proteins used in the cell-surface ACE2 staining experiments shown in Figure 2a. **b**, SARS-CoV-2 RBD in complex with human ACE2. The residues absent in donkey ACE2, as shown in Extended Data Figure 1, are indicated as magenta. **c**, Gating strategy for the flow cytometry data shown in Figures 2a and b. The left panel shows the preliminary FSC/SSC gate of the starting cell population. The right panel shows the gates that defines “negative” (red) and “positive” (purple) staining cell populations, respectively.

**Extended Data Fig. 3.**
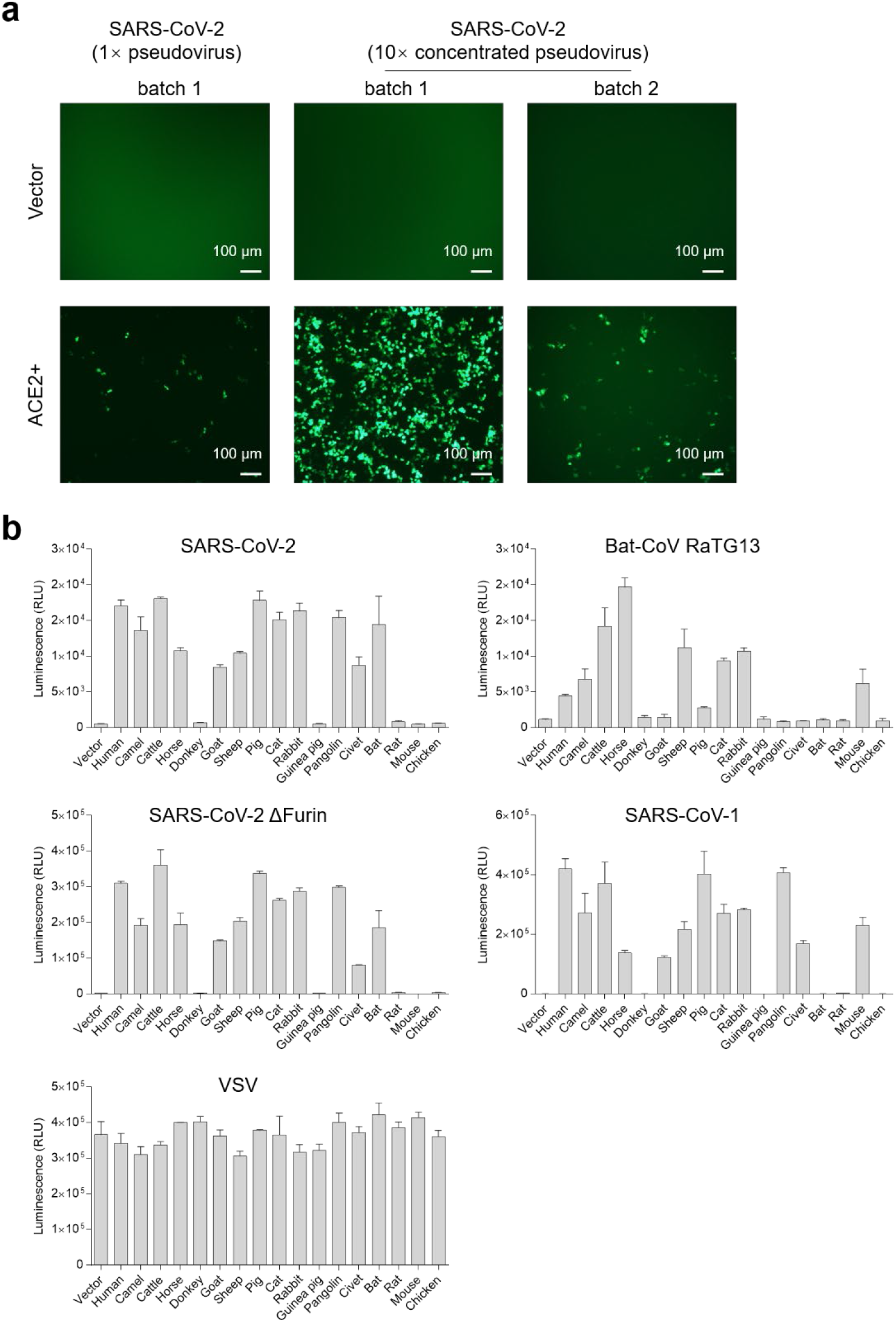
Optimization of the coronavirus pseudotype systems. **a**, Fluorescent microscopic images showing that SARS-CoV-2 wildtype spike-based pseudovirus system is not very robust and should be optimized. **b**, Original reporter expression data used for generating the heatmaps shown in Figure 3c. Data shown are representative of three experiments independently performed by three different people with similar results, and data points represent mean ± s.d. of three biological replicates.

**Extended Data Fig. 4.**
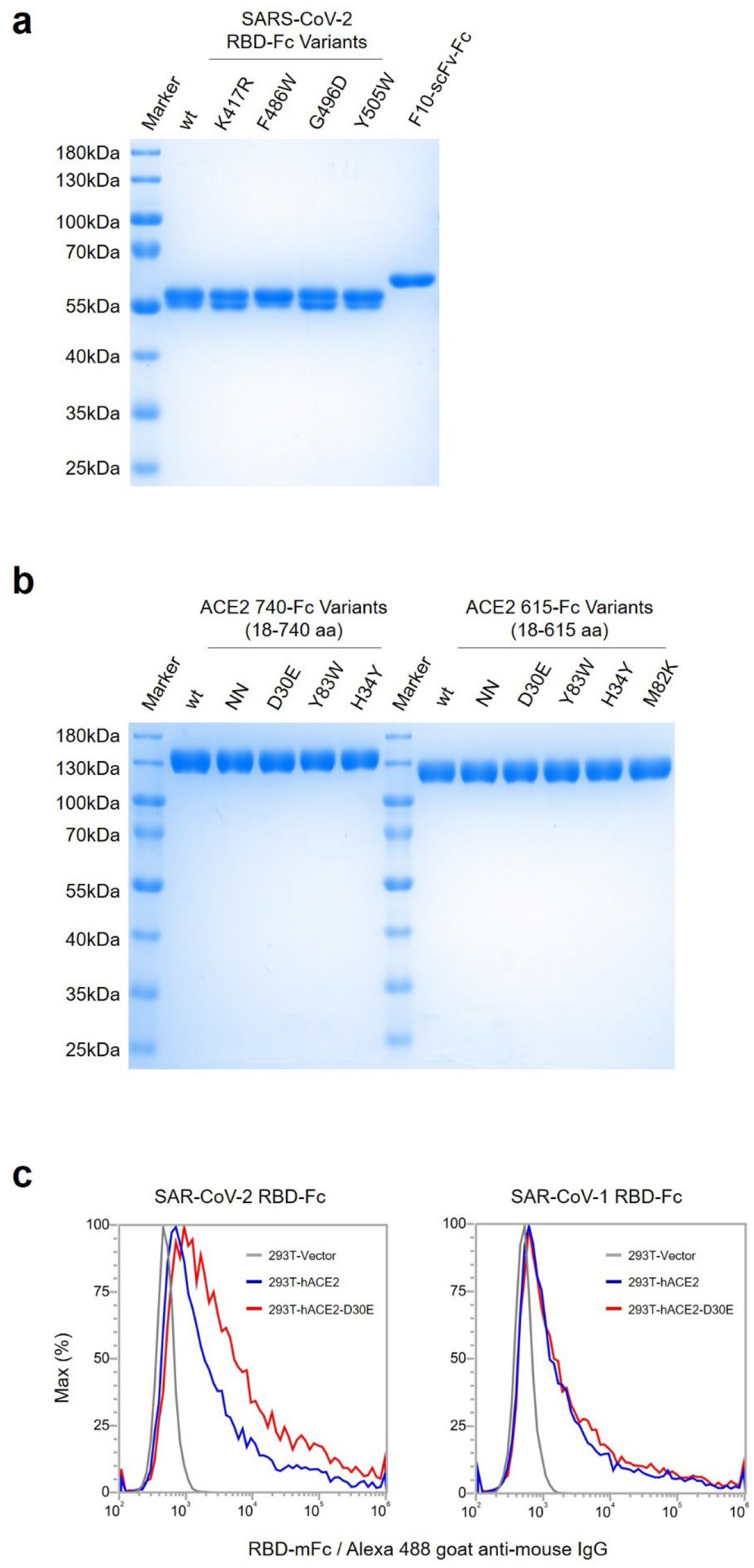
Key reagents (**a, b**) used in the experiments described in Figure 4 and flow cytometry data (**c**) supporting the D30E improvement of neutralization potency shown in Figure 4c.

**Extended Data Table 1.**
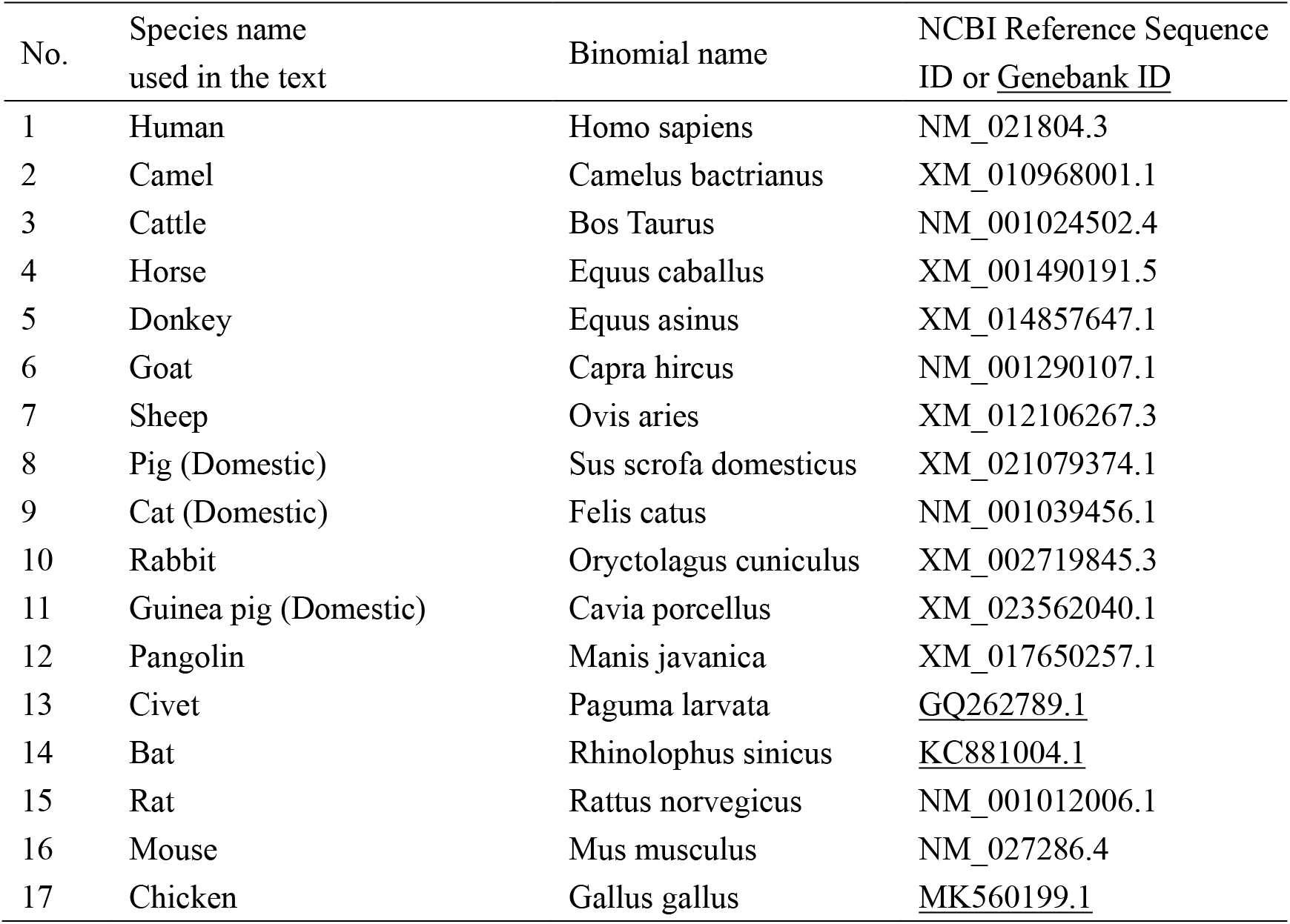
Species and accession numbers of ACE2 orthologs.

## Notes

### Competing Interest Statement

The authors have declared no competing interest.

